# Identification and characterization of novel infection associated transcripts in macrophages

**DOI:** 10.1101/2021.03.22.436546

**Authors:** A Prabhakar, Mohit Singla, Rakesh Lodha, Vivek Rao

**Author notes:** Address for correspondence: Vivek Rao.

## Abstract

Regulated expression of genes in response to internal and external stimuli is primarily responsible for the enormous plasticity and robustness of biological systems. Recent studies have elucidated complex regulatory non protein coding transcript (lncRNA) circuits in coordinated response of immune cells. By analysis of lncRNA expression profiles of macrophages in response to Mtb infection, we identified novel highly expressed transcripts, unique in encompassing one functional protein coding transcript- CMPK2 and a previously identified type I IFN responsive lncRNA- NRIR. While these RNA are induced by virulent Mtb early, the complete absence of expression in non-viable Mtb infected cells coupled to a more protracted expression profile in the case of BCG suggest an important role in macrophage response to mycobacteria. Moreover, enhanced expression was observed in macrophages from TB patients. The elevated expression by 1h in response to fast growing bacteria further emphasizes the importance of these RNAs in the macrophage infection response. These transcripts (TILT1, 2,3 - <u>T</u>LR4 and <u>I</u>nfection induced <u>L</u>ong <u>T</u>ranscript) are triggered exclusively by TLR4 stimulation (LPS) with faster and stronger kinetics in comparison to the lncRNA – NRIR. Overall, we provide evidence for the presence of numerous transcripts that is a part of the early infection response program of macrophages.

## Introduction

Intracellular bacteria encounter diverse host derived stresses and adapt to a wide ranging yet cell/ tissue specific niche(Huang and Brumell 2014; Hardison et al. 2018). Modulating the host to subdue these potentially harmful responses is one of the strategies employed by pathogens for long term infection (Cameron et al. 2015; Passalacqua et al. 2016). This plasticity requires an enormous degree of flexibility in the pathogen’s ability to not only sense the stimuli but fine tune host transcriptional programs to suit survival inside the cells (Jimenez et al. 2016; Cornejo et al. 2017; Eisenreich et al. 2017). It is not surprising that gene expression patterns at the onset of infection, immediate to the initial contact with host cells, often determine the outcome and overall pathogenesis of the organism.

Over several years, a detailed coverage of expression has helped identify protein coding patterns in diverse pathogens and cells/ intracellular niches in addition to paving the way for deciphering the regulatory circuits of non-protein coding transcripts(Winchell et al. 2016; Hayward et al. 2018). Recent evidences have delineated a definitive role for the long non-coding RNAs (lncRNA) in orchestrating the immune response by regulating gene expression is now being recognized (Imamura and Akimitsu 2014; Yu et al. 2015; Spurlock et al. 2016; Atianand et al. 2017). These RNAs have been implicated in the regulation of several cellular processes, immune cell functions and response to infections (Ouyang et al. 2014; Wang et al. 2014; Sigdel et al. 2015; Wang et al. 2015; Carpenter and Fitzgerald 2018; Xie et al. 2019; Menon and Hua 2020; Robinson et al. 2020). Non coding response dynamics have helped uncover novel facets of host – pathogen interactions in several bacterial pathogens (Gomez et al. 2013; Wang et al. 2015; Roy et al. 2018; Yan et al. 2018). Not surprisingly, Mtb infection of macrophages induces several of the non-coding transcripts with functional relevance in activating innate immune functions as well as modulation of these responses by the pathogen (Yang et al. 2016; Huang et al. 2018; Sharbati et al. 2019). Clinically relevant lncRNAs have recently been identified as putative biomarkers for TB patients (Chen et al. 2017; Li et al. 2017; Wu et al. 2020). With phagocytes initiating a strong innate immune response on primary contact with invading pathogens, it is logical to assume that activation of surface TLRs would be one of the initial signals for expression of gene regulatory networks in these cells. Recent advances in high throughput deep sequencing has revealed the identity of novel transcripts that arise either out of novel ORFS, trans-splicing events, long extensions and degradation products of previously identified transcripts (Srikumar et al. 2015; Kalam et al. 2017; Jackson et al. 2018; Agirre et al. 2019). In our attempt to study the role of type I IFN response in macrophages, we discovered that NRIR, previously identified as a lncRNA regulating this pathway was one of the highest expressed non coding transcript in Mtb infected macrophages. While the origins of these RNA is not yet clearly defined, detailed evaluation of this genomic locus indicated the presence of multiple transcripts. In line with this, we identified 3 other transcripts that appear to be encompassing NRIR and the neighboring IFN inducible gene – CMPK2. We demonstrate that these transcripts- TILT 1,2,3 are induced in response to bacterial infection temporally and is specific to activation of TLR4 by LPS. Expression of TILT is induced significantly in patients with active TB. Early expression following stimulation of macrophages suggests an important role for this transcript in the rapid response program of cells.

## Results

### The lncRNA NRIR is induced strongly in response to Mtb infection in macrophages

The response of macrophages to mycobacterial infection is characterized by an elaborate expression of genes that include protein coding and non-coding transcripts. The type I IFN signaling pathway is one of the early and robust response of host macrophage to mycobacterial infection (Novikov et al. 2011; Desvignes et al. 2012). Previous studies have identified the lncRNA NRIR as a primary lncRNA regulating this response in infected cells (Kambara et al. 2014). We hypothesized that NRIR, given the strong type I IFN signaling induced, would also be a crucial component of the host response to Mtb in macrophages. In line with our surmise, NRIR showed the highest expression levels in Mtb infected macrophages amongst a few of the well annotated mammalian lncRNAs (Fig. 1A). While NRIR was expressed 6h post infection with a steady increase in levels at later time points, the other innate response regulating lncRNA-NEAT1, was more protracted and was expressed at ~3x higher by 24h of infection (Fig. 1B). In fact, NRIR showed a steady rise in expression from the ~4x by 6h to a sharp increase by 24h of infection in excess of 200 times the values in uninfected samples (Fig.1C). In response to a non-pathogenic vaccine strain, BCG, NRIR was only induced by the later time point (24h) in infection, albeit at a lower magnitude than Mtb infection (Fig. 1D). Interestingly, the response was completely abrogated in macrophages exposed to heat killed Mtb even by 24h implicating an active infection mediated induction of these non-coding transcripts in macrophages. The kinetic response of NRIR induction observed in *S. typhimurium* and *E. coli* infections further validated the importance of these transcripts in infection response of human macrophage. NRIR was visible by the 1^st^ h of infection followed by a steady increase of expression by 6h of infection in macrophages infected by either strain of bacteria (Fig. 1E).

**Figure 1:**
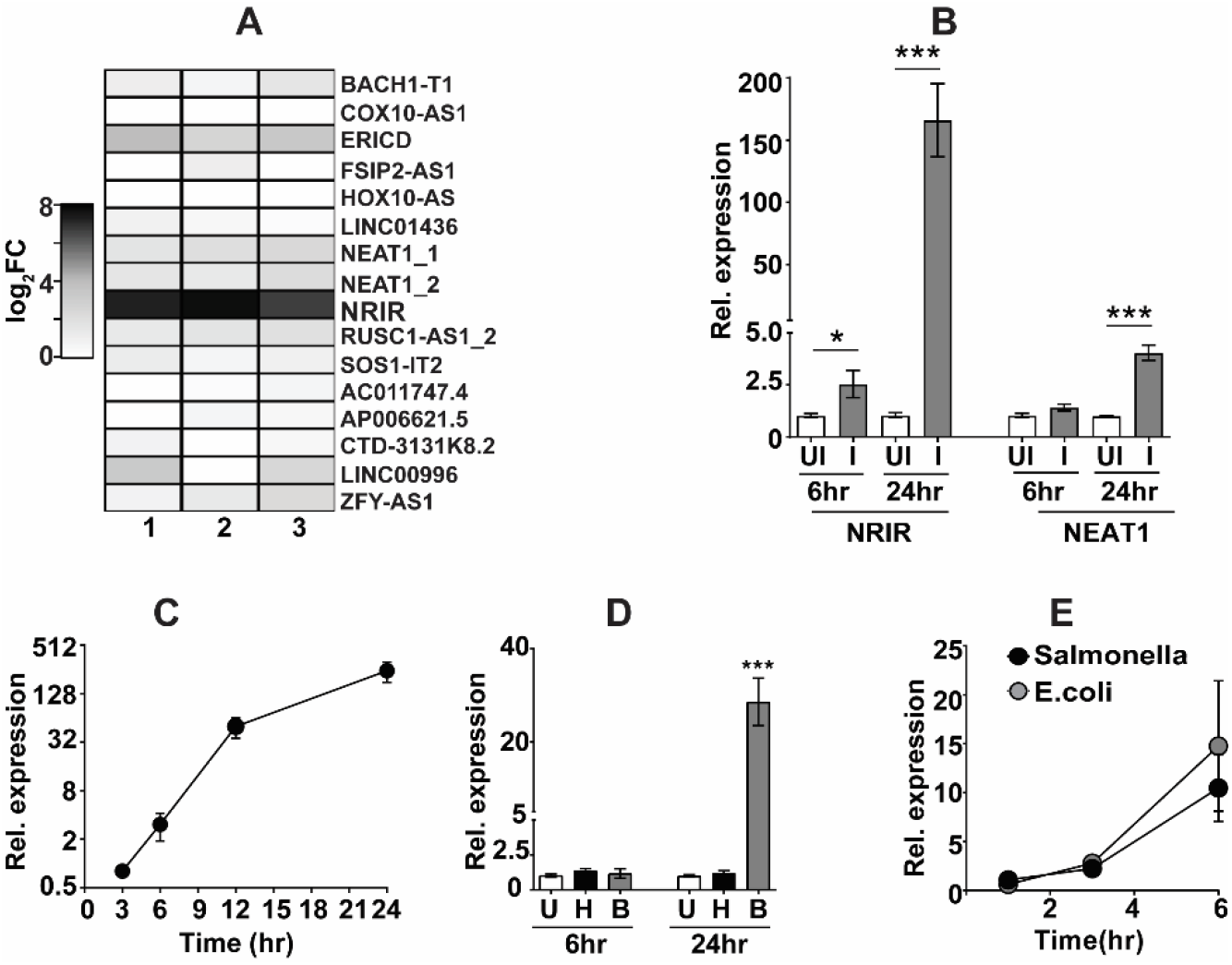
NRIR is expressed in response to bacterial infection. (A) The expression levels of a few of the annotated non-coding transcripts was analysed by qRT PCR in THP1 macrophages infected with Mtb. Change in expression of triplicate assays relative to GAPDH (log_2_FC) is depicted as a heat map. Three independent replicate experiments are individually represented as mean expression values from triplicate assays (n=3). B-E) Expression of NRIR (B-E) and Neat1 (B) in macrophages infected with Mtb (B-D) or *E. coli*/ or *S. typhimurium* (E) at different time intervals was analysed by qPCR. Change in expression of triplicate assays relative to GAPDH (logFC) is represented as rel. expression from triplicate assays of 2-3 independent replicate experiments. U- uninfected, H- Heat killed Mtb, B- *M.bovis* BCG.

### The NRIR locus supports transcription of several other non-coding transcripts

NRIR is localized in the same genomic loci with the mitochondria associated gene – CMPK2 (Fig. 2A). While NRIR is the most abundant lncRNA, there have been reports of the presence of other non-coding transcripts in this locus ((Lagarde et al. 2017), Fig.2A). To evaluate other infection specific transcripts, we used a PCR strategy with primers mapping to the NRIR/ CMPK2 specific region (P1 and P4) and observed three distinct amplicons of ~400bp, 800bp and 1kb which were absent when genomic DNA was used as a template (Fig. 2B). Both NRIR and CMPK2 specific amplicons were observed in reactions with the larger 2 fragments as template (Fig. 2C). Moreover, while the 800bp fragment produced the 400bp and 800bp amplicons, the largest 1kb fragment again yielded all 3 fragments on amplification with primers-1 and 4. Sequencing of the amplicons revealed the presence of 3 unique splice variants with fusion of Exon 4 of CMPK2-203 and the first exon of NRIR-202 (Fig. 2D). We then analyzed expression in total blood RNA of healthy individuals. As seen in THP1 cells, amplicons of ~800bp and 1kb in PCR with specific primers 1 and 4 were observed confirming the universal presence of these transcripts (Fig. 2E).

**Fig. 2.**
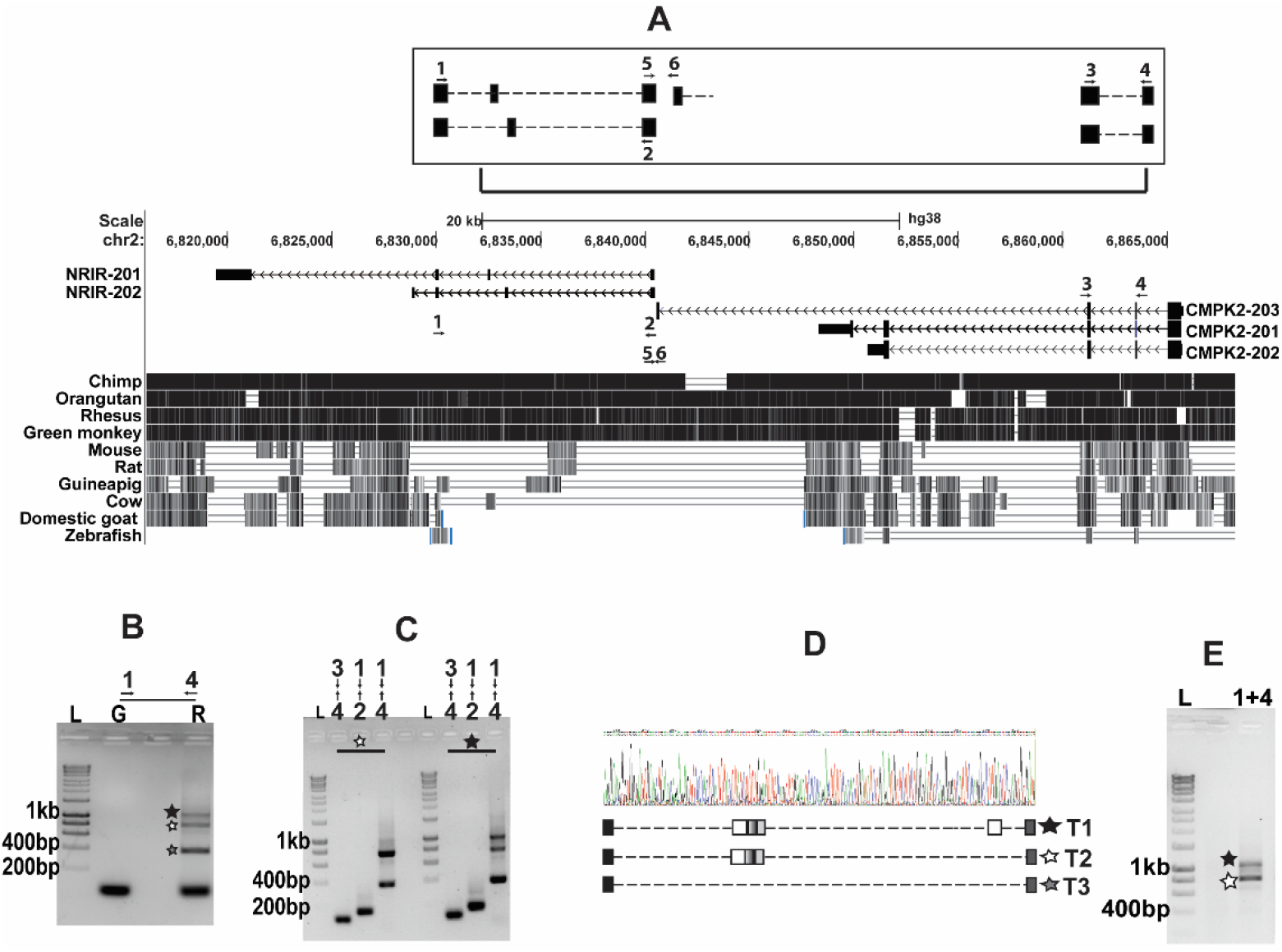
Identification of the novel transcripts-TILT encompassing CMPK2 and another lncRNA – NRIR in Mtb infected macrophages. A- The genomic locus of CMPK2 and NRIR in human cells is depicted with the previously annotated transcripts. The primers used for PCR analysis of transcripts are depicted by numbers. B, E- Analysis of the PCR amplification of cDNA (B, 24h) by using primers (1,4) from THP1 cells infected with Mtb as indicated to regions as depicted in panel A of the genomic locus by agarose gel electrophoresis. C- Amplification of the PCR product of reaction in B with primers as indicated in the figure. D- The profiles of exons present in the 3 new non-coding transcripts identified by sequencing of the PCR products. E- Expression of the novel transcripts in total blood of a healthy individual analysed by agarose gel electrophoresis.

### The novel transcripts are actively induced in response to bacterial infection of macrophages

Macrophages are typically designed to respond to assorted stimuli and alter their transcriptional profile to suit the incoming insult. Being a critical component of the host innate defense, macrophages are fine tuned for a rapid response to infections. TILT, in contrast to NRIR was more protracted and reduced- expression in macrophages was seen only by 12h of infection (4fold) with Mtb with a further increase to 14-fold by 24h (14-fold) (Fig. 3A). The absence of TILT expression in macrophages infected with heat inactivated Mtb even by 24h argued for the importance of active infection in the expression of TILT. Moreover, delayed expression of these RNAs only by 24h in response to infection by a non-pathogenic slow growing mycobacterial strain-*M. bovis* BCG further corroborated the importance in the response to pathogenic infection (inset). Clinical relevance in infection was highlighted by the significantly enhanced expression in the blood of active TB patients in comparison to healthy individuals (Fig. 3B). A similar profile of enhanced expression of TILT was observed for infection with Salmonella and *E. coli* (Fig. 3D), expression levels of TILT showed a strong peak (~25-40 fold) by 3h of infection followed by a rapid decline to around 10-fold greater than basal levels by 6h of infection associated with the faster growth rate of these bacteria in comparison slow growing mycobacteria.

**Fig. 3.**
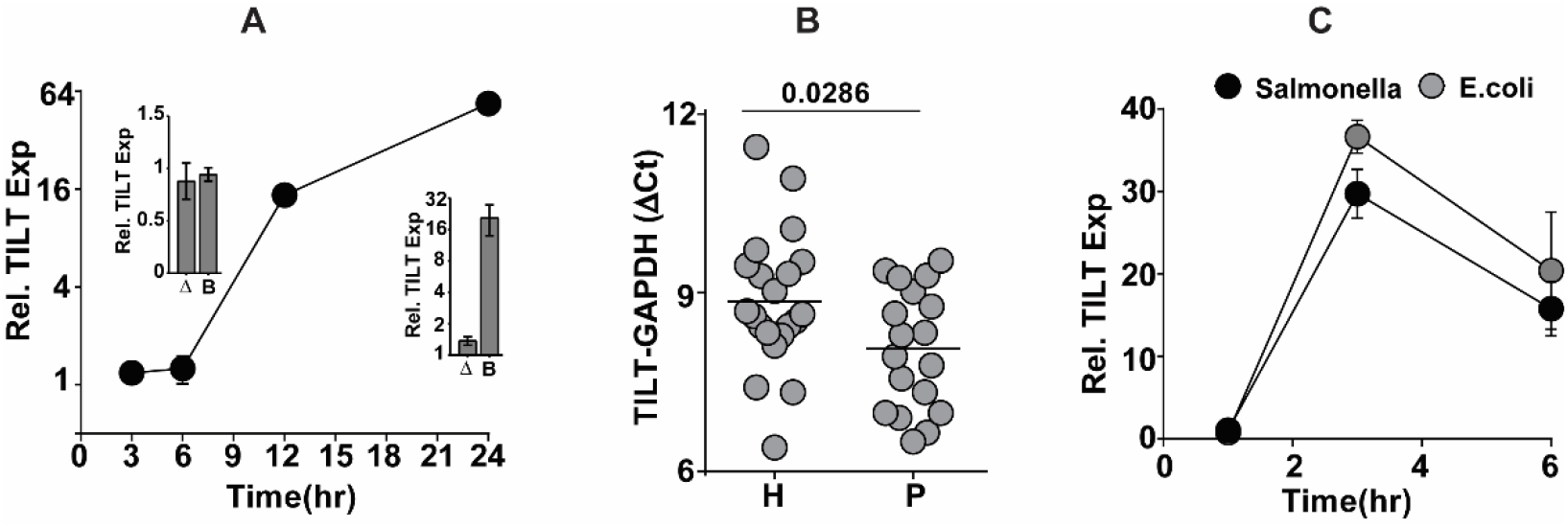
LncRNA-TILT is actively induced in Mtb infected macrophages. A-C, Expression kinetics of TILT in macrophages infected with Mtb at a MOI of 5. At the indicated intervals, RNA prepared from cells was used for analysis by qRT PCR. A- RNA from THP1 cells Inset- at 6 and 24h of infection with Heat killed Mtb (Δ) or BCG. B- from blood of healthy individuals and TB patients; each symbol represents one individual. C- in THP1 macrophages infected with *E. coli*, *S. typhimurium* at a MOI of 5 for the indicated time intervals. Values represented are mean + SEM of triplicate wells of 2-3 independent experiments.

### TLR4 signaling acts as the primary stimulus for TILT expression in macrophages

Macrophages, being the primary responders to any infection, are endowed with the capacity to recognize microbial PAMPS via innate receptors like TLRs and activate signaling mechanisms for rapid and robust neutralization of the pathogen. To identify the stimulus involved in activating this lncRNA, we analyzed TILT expression in macrophages stimulated with different TLR ligands (Fig. 4A). Where stimulation of most TLRs resulted in minimal induction, only LPS actively enhanced TILT expression in THP1 cells by ~ 20-fold increase by 6h and a sharp decline to ~9 fold by 24h. In response to LPS, the two transcripts- NRIR and TILT showed distinct kinetics of expression. Contrasting with the Mtb induced profile of faster NRIR expression, TILT was strongly induced as early as 3h (~16 fold) reaching to an excess of 200 times the basal levels by the 6^th^ hour of LPS stimulation and stabilizing thereafter to nearly 100 folds by 48h. In contrast, NRIR, despite any detectable expression by 3h, amplified rapidly to reach ~50 times basal levels with a peak of ~150 fold at 24h and then stabilizing to ~60 fold by 48h of stimulation (Fig. 4B). A complete dependence on TLR4 signaling was observed for expression of TILT in macrophages- treatment with the TLR4 antagonist -CLI095 completely neutralized TILT expression in macrophages in response to LPS, *E. coli* and *S. typhimurium*. In contrast, the effect of CLI0095 was only partial in THP1 infected with Mtb suggesting a TLR4 independent accessory signaling for TLT expression (Fig. 4C).

**Fig. 4.**
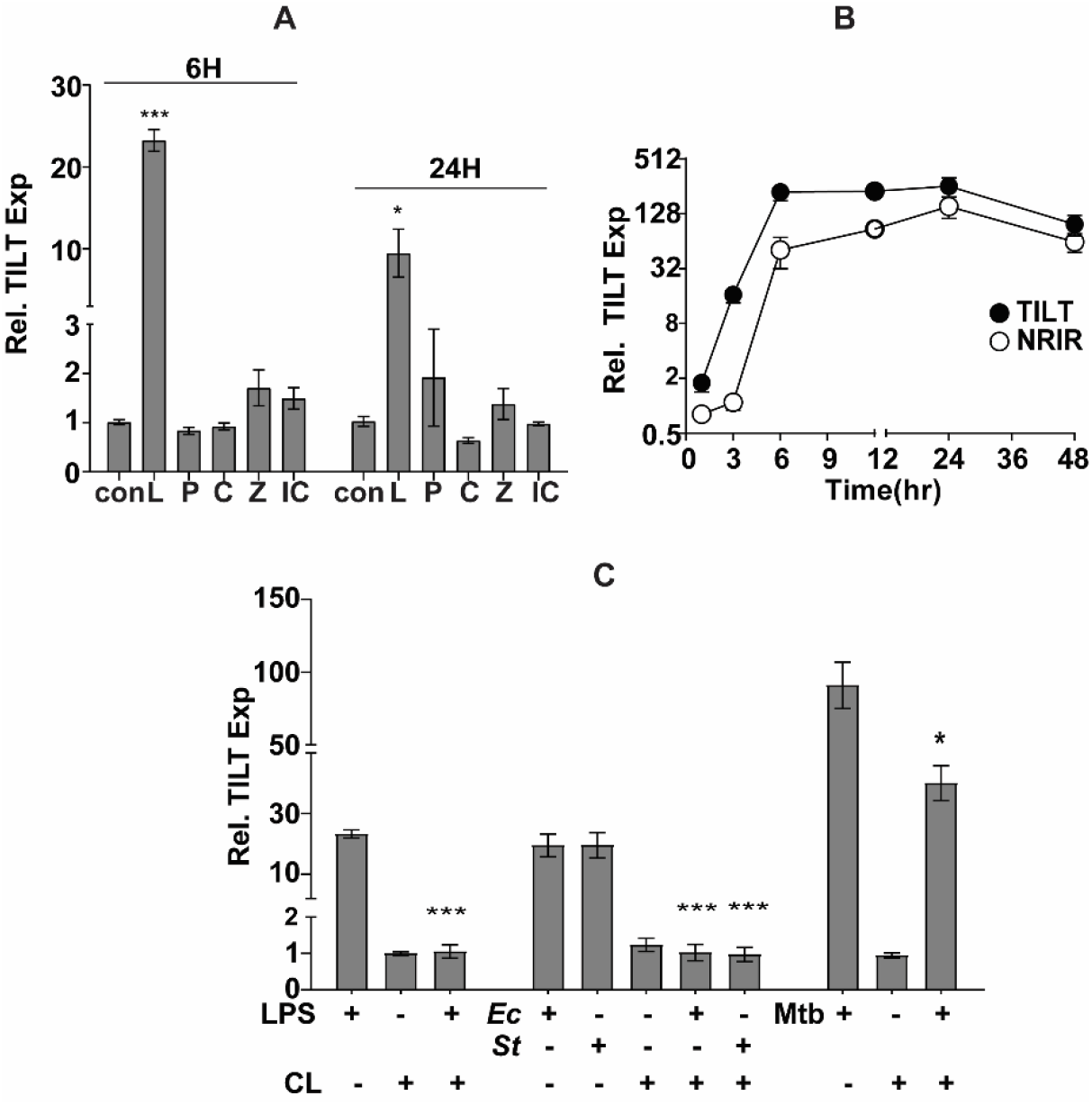
TILT is induced temporally in response to LPS stimulation in THP1 macrophages. A- TILT expression in macrophages stimulated with different TLR ligands- L- LPS at 10 ng/ml, P- Peptidoglycan at 20 ng/ml, C- Pam3CSK at 20 ng/ml, Z-Zymosan at 10 μg/ml, IC- poly IC at 2 μg/ml) at 6 and 24h post stimulation. B) Kinetics of TILT and NRIR expression in macrophages treated with 10ng/ml of LPS. C) TILT expression in macrophages infected with *S. typhi* (3h *p.i.*), *E. coli* (3h *p.i.*), Mtb (24h *p.i.*) or stimulated with LPS (6h *p.s.*) in the presence of the TLR4 inhibitor CLI095 (3 μM). At the indicated intervals RNA prepared from cells was used for analysis by qRT PCR and normalized to GAPDH. Relative expression at any given time w.r.t unstimulated is depicted as mean ± SEM for triplicate assays of 2-3 independent experiments.

## DISCUSSION

Macrophages have been shown to successfully respond to numerous stimuli including infections and initiate response profiles for control of the pathogen (Weiss and Schaible 2015). Over several decades of research, the landscape of stimuli driven protein coding expression patterns have been realized in high detail (Roy et al. 2018). It is now possible to portray the layout of protein/ pathway induction profiles in host macrophages as a response to infection with several bacterial pathogens (Denzer et al. 2020). The discovery of several non-protein coding transcripts as key regulators has only supplemented another layer to the multi-scale pyramid of the macrophage response kinetics (Aune and Spurlock 2016; Duval et al. 2017; Zur Bruegge et al. 2017; Ma et al. 2019; Ahmad et al. 2020). In the last few years, several non-coding transcripts and regulatory networks have been identified with important regulatory roles in infection with the help of powerful omics-untargeted / specific-targeted approaches (Moon et al. 2014; Liu et al. 2018; Menard et al. 2018; Fan et al. 2019; Liu et al. 2019).

By using simple molecular techniques, we have identified novel non coding transcripts (TILT1-3) encompassing a previously identified lncRNA (NRIR) and a mitochondrial resident gene (CMPK2). Recently, by using high throughput Capture Long Seq (CLS), GENCODE consortium have re annotated lncRNAs population in human and mouse tissues identifying several noncoding transcripts (Lagarde, 2017 #77}. We demonstrate the presence of multiple transcripts comprising of different exons from the NRIR and CMPK2 genomic locus implying a higher degree of splicing based regulation of gene expression in this region. Our finding of TILT in human blood cells of a normal individual also argues for the presence of these transcripts in myeloid cell populations. Surprisingly, this arrangement of NRIR with CMPK2 is only observed in human cells and is absent from murine cells suggestive of a unique regulatory network to fine tune macrophage response kinetics (Breschi et al. 2017).

Despite a differential amplitude and kinetics of expression of the two non-coding transcripts (NRIR and TILT) in response to different stimuli, the rapid and sustained levels of expression over 24-48h of stimulus highlights the importance of this locus in the response kinetics of macrophages. Increased expression of TILT in sera of active TB patients only signifies the importance of this response in TB infections, coupled to the enhanced expression of TILT in response to infection with *E. coli* and Salmonella argues for an important regulatory circuit in macrophage response to infection. Further the strong induction of TILT with LPS stimulation earlier than the expression of NRIR is suggestive of a differential control of expression. While we did not find promoter like element different from the annotated promoter of CMPK2 (Data not shown), separate promoter like elements were observed upstream of NRIR indicative of an uncoupled expression of this lncRNA from CMPK2 and TILT. Abrogation of TILT expression following treatment with CLI095 in LPS stimulated, in *E. coli*, Salmonella infections only substantiates the absolute requirement of TLR4 mediated signaling for TILT expression in macrophages.

In sharp contrast to NRIR, expression of TILT was more protracted and reduced in comparison in response to Mtb infection. Moreover, treatment of Mtb infected cells with CLI095 only partially reduced the expression of TILT in macrophages. It is well established that macrophage response to Mtb is not primarily dependent on TLR4 signaling with combinatorial requirement for other TLR (TLR2), NLR, RLR mediated signaling rather than predominant signaling axis (Kleinnijenhuis et al. 2011; Faridgohar and Nikoueinejad 2017). In fact, mice lacking multiple TLR did not suffer from increased susceptibility to Mtb infection (Nguyen et al. 2020) again corroborating the multiple axis of macrophage stimulation by Mtb leading to the enormous redundancy and plasticity of the macrophage responses. Identifying the regulatory circuit and the consequence of this control in macrophage/ immune cell function would provide important insights into the diversity and complexity of mammalian innate responses.

## Material and Methods

### Bacterial Strains and Growth Conditions

*Mycobacterium tuberculosis* strain Erdman was grown in 7H9 Middlebrook enriched with Middlebrook ADC (BD Biosciences, USA). *E. coli* and *Salmonella typhimurium* were grown in LB media (BD Biosciences, USA) at 37°C.

### Macrophage culture and Infection

RPMI 1640-GlutaMAX (Himedia laboratories, Mumbai, India) supplemented with 10% fetal bovine serum (Himedia laboratories, Mumbai, India,) and 1mM sodium pyruvate (Himedia laboratories, Mumbai, India) was used to culture THP-1 monocytes. Differentiation of THP1 monocytes to macrophages was done using 100nM phorbol myristate acetate (PMA) (Sigma Aldrich, USA). Mtb, *E. coli* and *Salmonella typhimurium* were grown to mid-log phase at 37°C, washed twice with phosphate buffered saline (PBS) containing 0.05% Tween80, finally suspended in PBS and centrifuged at 800 rpm for 10 min to get a uniform single cell suspension. This uniform single cell suspension was diluted with complete RPMI 1640 to the required cell density and then used to infect the differentiated THP-1 macrophages at multiplicity of infection (MOI) of 5. At different time intervals, cells were harvested in RNAzol (Sigma Aldrich, USA) for analysis. LPS, CLI095, poly I:C, and Pam3CSK2 were purchased from (InvivoGen, Toulouse, France). Zymosan (Sigma Aldrich, USA) was used according to the concentration mentioned.

### RNA isolation and qRT-PCR

RNA isolation was done as per standard RNAzol protocol. cDNA synthesis was performed from 1μg RNA using verso cDNA synthesis kit (Thermo Scientific, USA). The expression level was checked by DyNAmo Flash SYBR Green qPCR kit (Thermo Scientific, USA) in Roche LC480 system. *Gapdh* was used as internal control. Ct values were normalized with uninfected control.

### Genome locus and PCR

Genome locus was analyzed using UCSC web (https://genome.ucsc.edu/.)

### Whole blood collection

Confirmed cases of pulmonary TB patients between ages of 5 and 15 years were included in the study with prior consent as per the Institutional ethical committee guidelines. BCG vaccination, primary or secondary infection and single or co-infection status were recorded. HIV positive and any non-respiratory major disease patients were excluded from the study. The control samples were obtained from individuals without any overt clinical manifestation of disease. 2-3 ml blood was collected in RNAgard blood tubes (Biomatrica, USA) and stored in −80°C until RNA precipitation. RNA was isolated from frozen blood samples as per the manufacturer’s recommendations.

### Graphs

Graphs were generated using Graphpad prism or R. Student t test was used for statistical analysis of the data.

**Table 1:**
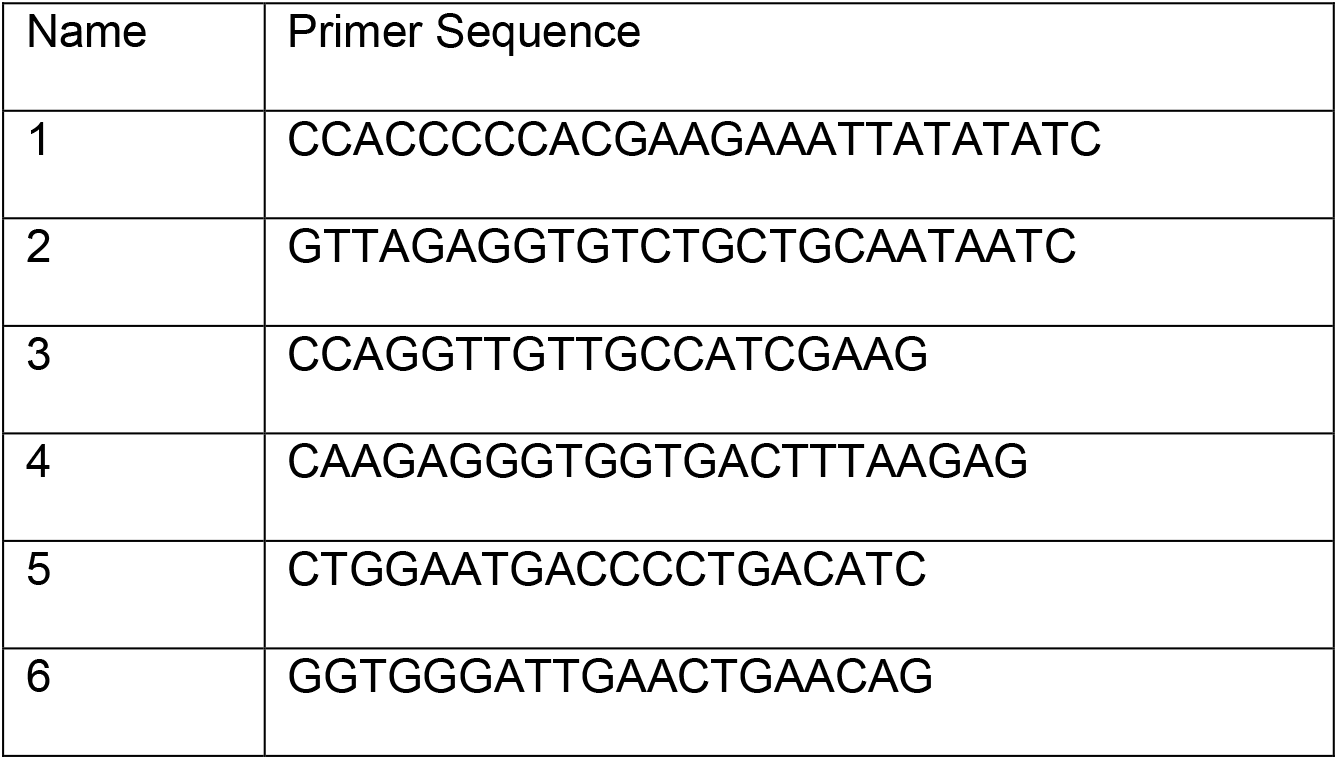
List of primers used in the study.

## Abbreviations

IFN: Interferon
NRIR: 
CMPK2: 

## Acknowledgements

The authors thank CSIR (VR-BSC0123) and STS0016 for funding and facility maintenance and PA-CSIR-BSC0124, CSIR SRF/ RA for fellowship.

## Author Contributions

PA, VR were involved in conceptualizing and design of the work, MS and RL were instrumental in sampling of blood from TB patients, the work was performed by PA and manuscript was written by VR and PA.

